# Modeling the frequency and number of persons to test to detect and control COVID-19 outbreaks in congregate settings

**DOI:** 10.1101/2020.11.20.391011

**Authors:** Prabasaj Paul, Emily Mosites, Rebecca L. Laws, Heather Scobie, Rachel B. Slayton, Kristie E. N. Clarke

## Abstract

**Background:** Congregate settings are at risk for coronavirus disease 2019 (COVID-19) outbreaks. Diagnostic testing can be used as a tool in these settings to identify outbreaks and to control transmission.

**Methods:** We used transmission modeling to estimate the minimum number of persons to test and the optimal frequency to detect small outbreaks of COVID-19 in a congregate facility. We also estimated the frequency of testing needed to interrupt transmission within a facility.

**Results:** The number of people to test and frequency of testing needed depended on turnaround time, facility size, and test characteristics. Parameters are calculated for a variety of scenarios. In a facility of 100 people, 26 randomly selected individuals would need to be tested at least every 6 days to identify a true underlying prevalence of at least 5%, with test sensitivity of 85%, and greater than 95% outbreak detection sensitivity. Disease transmission could be interrupted with universal, facility-wide testing with rapid turnaround every three days.

**Conclusions:** Testing a subset of individuals in congregate settings can improve early detection of small outbreaks of COVID-19. Frequent universal diagnostic testing can be used to interrupt transmission within a facility, but its efficacy is reliant on rapid turnaround of results for isolation of infected individuals.

## Background

SARS-CoV-2, the virus that causes coronavirus disease 2019 (COVID-19), is a highly transmissible pathogen that spreads easily in shared environments (1). Thus, congregate settings, such as long-term care facilities, correctional facilities, homeless shelters, and high-density workplaces, are at increased risk for outbreaks of COVID-19 (2-9). Diagnostic testing to detect SARS-CoV-2 infection in congregate settings may achieve at least two key public health objectives: 1) testing can identify outbreaks, triggering the application of intervention measures and further testing, and 2) testing can identify infectious individuals who need to isolate to prevent further transmission.

We developed and applied models related to SARS-CoV-2 infectiousness, test sensitivity and specificity, and time to test results to estimate both the number of people and frequency of testing needed in congregate settings to achieve: 1) early detection of outbreaks and 2) transmission interruption. These estimates can inform testing strategies designed to protect people living or working in congregate settings and the communities in which they reside.

## Materials and Methods

### Testing for early detection of outbreaks

Outbreak detection was defined as identifying at least one positive test, regardless of true underlying prevalence. We defined “early” detection to mean identifying at least one positive result before case counts surpass a set number; here we chose 2, 5, and 10 true cases. For this purpose, the number of people that need to be tested in a congregate setting is dependent on the true underlying prevalence of infection, and the sensitivity and the specificity of the test. We estimated the number of randomly-selected individuals (*n*) needed to test for detection of an outbreak based on an expected number of infections (*n*_0_, commensurate with prevalence *p*), a minimum detection sensitivity *S*, and a minimum positive predictive value. When the *n* required exceeded the facility size *N*, then the outbreak was considered too small to be detected.

If *n* individuals are randomly selected for testing (test sensitivity ϵ, specificity π) from a facility of size *N* where the prevalence of infections is *p*, and presence of an outbreak is indicated by at least one positive test result, the probabilities of four possible outcomes are:

- Detection of outbreak and an outbreak exists (true positive): *TP* = 1 − (*FP* + *FN* + *TN*)
- Detection of an outbreak but no outbreak exists (false positive): *FP* = (1 − *p*)^*N*^(1 − π^*n*^)
- No detection of an outbreak but outbreak exists (false negative): *FN* = [*p* (1 − ϵ) + (1 − *p*)π]^*n*^

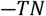
- No detection of an outbreak and no outbreak exists (true negative): *TN* = (1 − *p*)^*N*^π^*n*^

For the facility level, outbreak detection sensitivity is *S* = *TP* /(*TP* + *FN*) and positive predictive value is *PP V* = *TP* /(*TP* + *FP*).

The true underlying prevalence of infection, *n*, was determined by the rate of introduction, (η = community incidence × facility size) and the doubling time (τ) within the facility as:

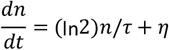

Solving for the expected number of infections at time *t* after first introduction (i.e., with n(0)=1) gives:

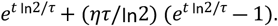

At a facility where the rate of introduction is η and the doubling time is τ, the expected number of infections at time *t* after first introduction is:

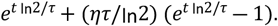

We assumed a community incidence of 100 cases per 100,000 people and doubling time for congregate settings of 3.4 days (4). Expressions were calculated using R software v3.6.2.

### Testing to interrupt transmission

To interrupt transmission (i.e., achieve a basic reproduction number below one), a successful testing strategy may need to achieve at least 60% reduction in transmission (10). We estimated the percent reduction in transmission for two scenarios of facility-wide testing: 1) both test-positive and symptomatic individuals are isolated, and 2) only test-positive individuals are isolated (appropriate in settings where symptom ascertainment is difficult). Expressions were also calculated using R 3.6.2.

We used the infectivity profile of COVID-19 estimated by He *et al*. (11). We also used an assumption that presymptomatic infections account for 50% of transmission and 40% of infections are asymptomatic (12). In our model, test sensitivity ranged 85%–95%, depending on both the test type and time since exposure (i.e., more sensitive during times of high viral load) (13).

If the infectiousness of an infected individual is *I*(*t*) (normalized to unity), where *t* is the time since exposure, and the fraction of infected individuals who have already shown symptoms at *t* is σ(*t*), then the proportion of transmissions eliminated on isolation of symptomatic individuals, *P*_*s*_, is:

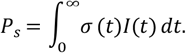

If testing is repeated on the same individual at regular intervals *T*, then the mean proportion of asymptomatic transmissions eliminated is:

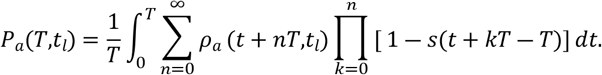

If an asymptomatic infected individual is tested at time *t*_0_ after infection, with test sensitivity *s*(*t*_0_), and the test result is available after a reporting lag *t*_*l*_ at *t*_1_ = *t*_0_ + *t*_*l*_, the additional proportion of transmissions eliminated, *ρ*_*a*_, is:

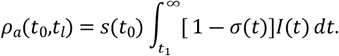

If testing is repeated on the same individual at regular intervals *T*, then the mean proportion of asymptomatic transmissions eliminated is:

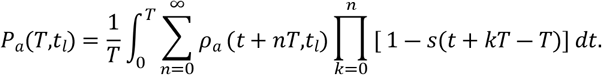

The sum accounts for those who tested true positive when first tested, as well as those who tested false negative once, twice, etc. followed by a true positive.

## Results

### Testing for early detection of outbreaks

We estimated the number of individuals needed to test for early outbreak identification for 18 scenarios (Table 1), which required testing between 16% and 90% of individuals, depending on facility size and true cluster size. For a facility of 100 people, 47 individuals would need to be tested to identify at least one positive result if there were truly 5 cases (5% prevalence) with test sensitivity of 85%, and greater than 95% outbreak detection sensitivity.

**Table 1:** Number of individuals needed to test to detect at least one positive result at facilities of varying sizes given different numbers of cases present (test sensitivity 85%, detection sensitivity ≥ 95%)

| Facility Size | Number of cases at facility |  |  |
| --- | --- | --- | --- |
|  | 2 cases | 5 cases | 10 cases |
| Number of individuals needed to test |  |  |  |
| 30 | 27 | 17 | 9 |
| 50 | 42 | 27 | 15 |
| 100 | 69* | 47 | 28 |
| 150 | 86* | 61 | 38 |
| 300 | 110* | 87 | 61 |
| 500 | 123* | 104 | 80 |

\*Positive predictive value ≤ 90%

The frequency of testing for detection at a facility should not exceed the time from first introduction of an infected person from the community to the maximum threshold of allowable infections, *n*_0_. At a 100-person facility located in a community with an incidence of 100 infections per 100,000 daily, an introduction from the community can be expected to occur on average every 10 days, and a 5% prevalence would be attained 6 days after introduction (Table 2). Therefore, the interval between tests should not exceed 6 days to detect an outbreak at 5% prevalence or lower.

**Table 2:** Average number of days—from first introduction of an infected individual from the community—to reach a given number of infections at facilities of varying sizes

| Facility Size* | Average number of days between introductions** | Days until x number of cases reached |  |  |
| --- | --- | --- | --- | --- |
|  |  | 2 cases | 5 cases | 10 cases |
| 30 | 33 days | 3 days | 7 days | 11 days |
| 50 | 20 days | 3 days | 7 days | 10 days |
| 100 | 10 days | 3 days | 6 days | 10 days |
| 150 | 7 days | 2 days | 6 days | 9 days |
| 300 | 3 days | 2 days | 5 days | 8 days |
| 500 | 2 days | 1 day | 4 days | 6 days |

\*Doubling time is 3.4 days within the facility regardless of size.
\*\*If true community incidence is 100 infections daily per 100,000, this is the average number of days in between introduction of infected individuals to the facility, given the facility size.

### Testing to interrupt transmission

With no testing, isolation of symptomatic individuals at symptom onset alone would achieve a 33% reduction in transmission (Figure 1A). When symptom-based isolation and testing with immediate results (within 15 minutes) were combined, a 60% reduction in transmission (i.e., R_0_ <1) (1) would be achieved if tests were administered at least every 3 days. A 60% reduction in transmission would also be achieved by administering tests every 2 days if there was a 24-hour delay in results reporting, and by administering tests daily with a 36-hour delay in results.

**Figure 1.**
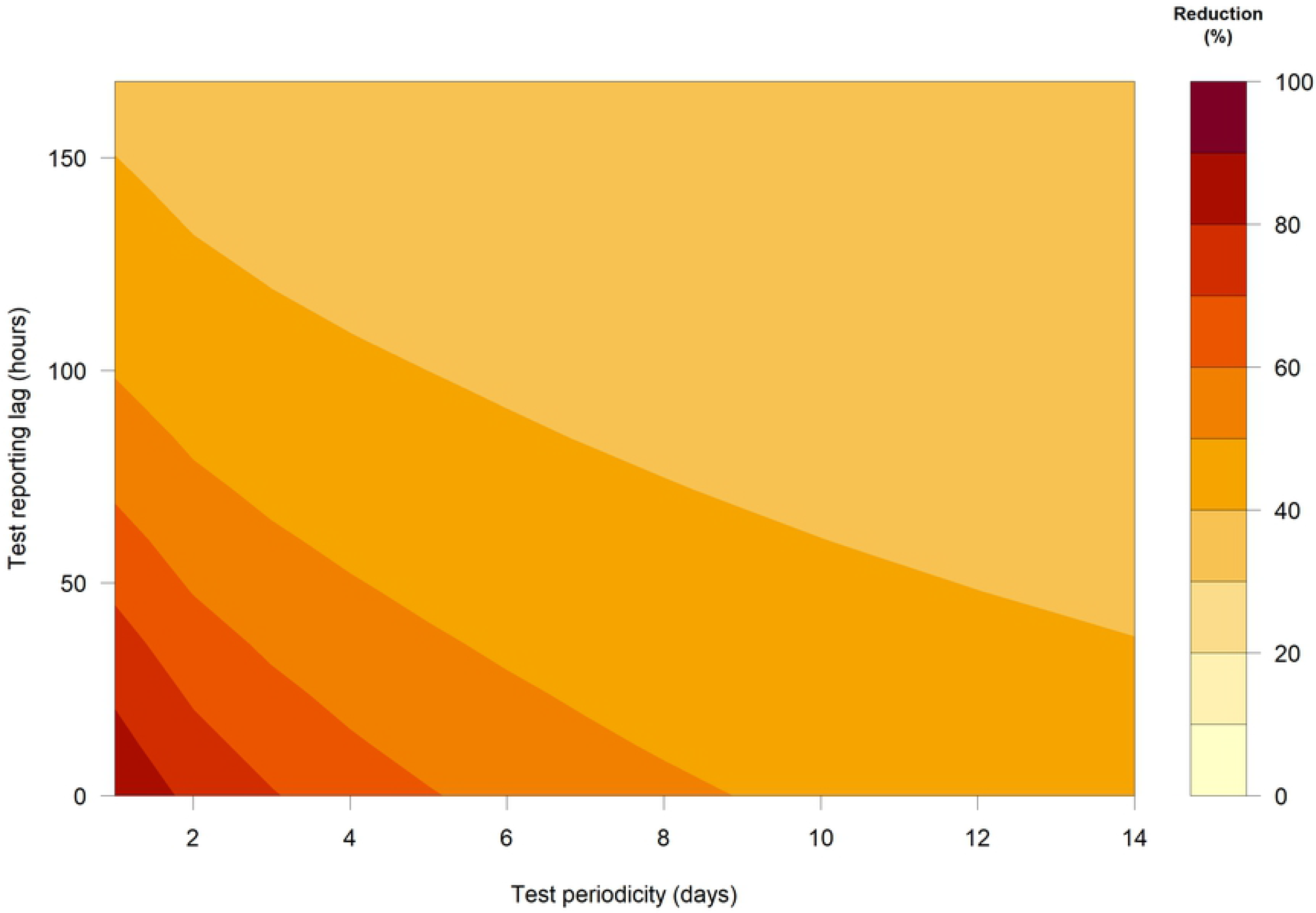
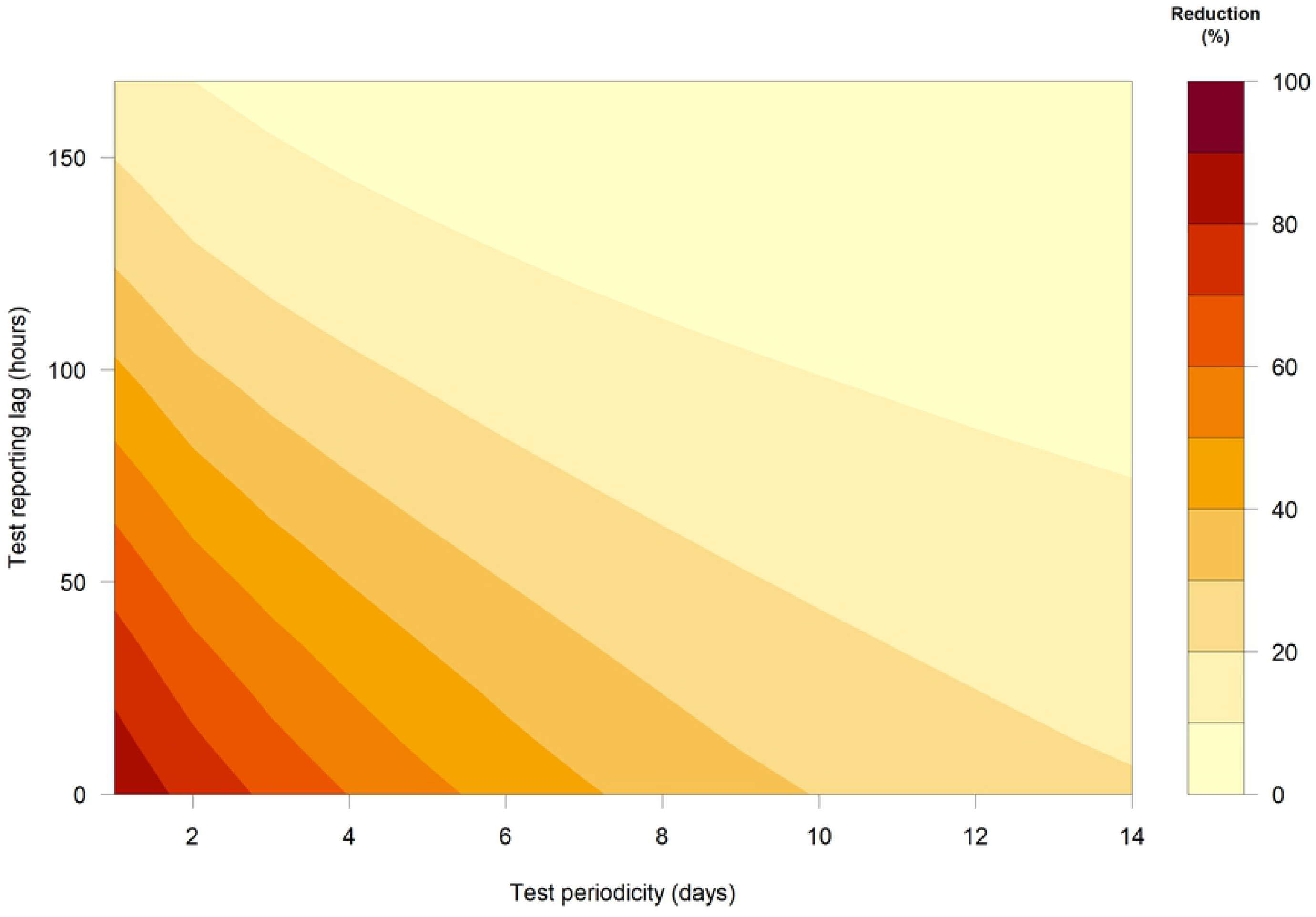
(A) Scenario 1, Reduction in transmission (%) through isolation of test-positive and symptomatic individuals. (B) Scenario 2, Reduction in transmission (%) through isolation of test-positive individuals only.

Using testing alone without additional isolation of symptomatic individuals (Figure 1B) would require more frequent testing to achieve a 60% reduction in transmission. Daily testing would be required if results were available in 24 hours and testing every 2 days required if there was a 12-hour delay in test results.

## Discussion

Numerous COVID-19 outbreaks have occurred in congregate settings, sometimes with high morbidity and mortality (2-9). In this modeling study, we found that early identification of an outbreak of 5 cases in a facility could be achieved by testing from 21% to 57% of individuals, depending on facility size. Minimal testing intervals were estimated (e.g., every 6 days for a 100 person facility), but more frequent testing would increase the likelihood of early detection. Testing frequency should take into account facility size. Detection of cases among a sample of the facility population would indicate a need for facility-wide testing and other intervention measures to interrupt transmission.

Using testing as a tool to interrupt transmission required a much higher frequency of testing, generally every 3 days or fewer, particularly if there are reporting lags in receiving test results (i.e., 1–3 days vs. 15 minutes). Findings regarding testing frequency align with another recent modeling study focused on institutes of higher education (14). Ultimately, the frequency of testing at a facility will depend on the balance between risk tolerance, frequency of introducing infections, and resource availability.

One limitation of this analysis is that we applied general formulas that did not account for specific characteristics of individuals residing in and working in these congregate settings. Furthermore, uncertainties in the modeling parameters introduce imprecision in derived estimates. These values can provide starting points for consideration for testing strategies but should not be considered definitive. As additional data become available, parameter estimates can be refined and models can be fitted to the best available data.

To reduce transmission, testing should always be used in combination with other prevention measures including social distancing, wearing masks and cloth face coverings, hand hygiene, cleaning and disinfection, screening, and isolation of individuals that are symptomatic or test positive, and quarantine of their close contacts. In coordination with these measures, testing strategies may reduce morbidity and mortality among individuals in congregate settings, prevent further spread into the community, and decrease strain on healthcare systems.

## Acknowledgments

The authors would like to thank Dr. Eric Mooring for his thoughtful review of the manuscript.

